# Differential effect of low and high frequency transcranial magnetic stimulation on cortical excitability and myelination after neonatal hypoxia in mice

**DOI:** 10.1101/2025.06.13.659545

**Authors:** Rafael Bandeira Fabres, Ivan Goussakov, Sylvia Synowiec, Emilia Schwenk, Vasiliy Yarnykh, Daniil Aksenov, Alexander Drobyshevsky

## Abstract

Premature infants are highly prone to intermittent hypoxic brain injury, which is linked to adverse motor, cognitive, and behavioral outcomes, including attention deficits, hyperactivity, and learning difficulties. Previous animal studies have revealed myelination deficits and increased glutamatergic synaptic strength in the sensory-motor cortex. This study examines the feasibility, safety, and therapeutic potential of repetitive transcranial magnetic stimulation (rTMS) to improve central hypomyelination, reduce excessive glutamatergic transmission in cortical neurons after neonatal intermittent hypoxia (IH), and enhance behavioral outcomes. In a mouse model of neonatal IH brain injury, low-frequency (LF-rTMS) at 1 Hz or high-frequency (HF-rTMS) at 10 Hz was administered for 5 days shortly after the injury. The rTMS treatment did not cause apoptosis or inflammation. HF-rTMS notably ameliorated hypomyelination in the corticospinal tract in both the stimulated and non-stimulated hemispheres. LF-rTMS decreased hyperactivity in female IH mice and lowered the heightened glutamatergic synaptic excitability in motor cortex slices. The data suggest that rTMS can affect both myelination and synaptic excitability, leading to improved behavioral outcomes following neonatal hypoxic brain injury. These results support the potential of rTMS as an early intervention for neurological issues caused by perinatal hypoxia.

## INTRODUCTION

Premature infants are highly susceptible to hypoxic brain injury, which often results in adverse neurological outcomes. A leading cause of motor deficits after perinatal hypoxic brain injury, including cerebral palsy, is disrupted ascending and descending motor control due to white matter injury (WMI) that affects axons and myelin sheaths ^1, 2^, especially in the posterior limb of the internal capsule (IC), where corticospinal tract (CST) fibers descend^3^. Although ongoing efforts aim to reduce cellular damage after brain injury, treatment options for established motor deficits are mainly palliative, with many therapies for spasticity and motor control issues focusing on symptom relief rather than addressing the root causes^4-6^. Restoring myelination and normal functional connectivity within motor control pathways would target the underlying source of motor deficits. Ideally, this should be done early to prevent the formation of abnormal circuits in infants with perinatal brain injury ^7, 8^.

Early interventions that stimulate motor control can be enhanced with non-invasive brain stimulation (NIBS), specifically repetitive transcranial magnetic stimulation (rTMS)^9^, to promote functional connectivity in motor control pathways. NIBS has demonstrated a high margin of safety in pediatric settings ^10, 11^. While showing promise in treating pediatric neurological conditions like stroke, epilepsy, and unilateral cerebral palsy ^12^, its potential for promoting myelination early after perinatal brain injury requires further exploration. The current study investigates the feasibility and effectiveness of rTMS in promoting activity-dependent myelination in corticospinal and other major white matter tracts.

In addition to motor abnormalities, adverse cognitive outcomes, including deficits in memory, attention, and executive function, are common after perinatal hypoxic brain injury ^13-17^. Animal data indicate that hyperactivity and learning deficits following neonatal intermittent hypoxia (IH) are linked to an abnormal increase in glutamatergic synaptic strength (synaptic potentiation), resulting in the occlusion of synaptic plasticity, a process crucial for brain development and cortical network formation ^18^. This underscores the need for therapeutic interventions that can target and, if possible, restore normal synaptic function.

While pharmacological methods to reduce excessive glutamatergic transmission face limitations due to the potentially harmful effects of long-term global inhibition on the developing brain, non-invasive neuromodulation offers a valuable alternative. However, the long-term effects and optimal stimulation settings of NIBS on the developing brain remain uncertain. This study examines the distinct effects of low- and high-frequency repetitive transcranial magnetic stimulation (rTMS) on cortical excitability and myelination in mice following neonatal hypoxic brain injury. We hypothesize that low-frequency rTMS can reduce excessive glutamatergic transmission in cortical neurons following neonatal intermittent hypoxia (IH), thereby helping restore synaptic plasticity, balance excitation and inhibition, and improve behavioral outcomes. Additionally, we aim to evaluate the feasibility and effectiveness of rTMS in enhancing myelination of the CST.

## METHODS

The study has received approval from the Institutional Animal Care and Use Committee of Endeavor Health. The experimental timeline is shown in Figure 1.

**Figure 1.**
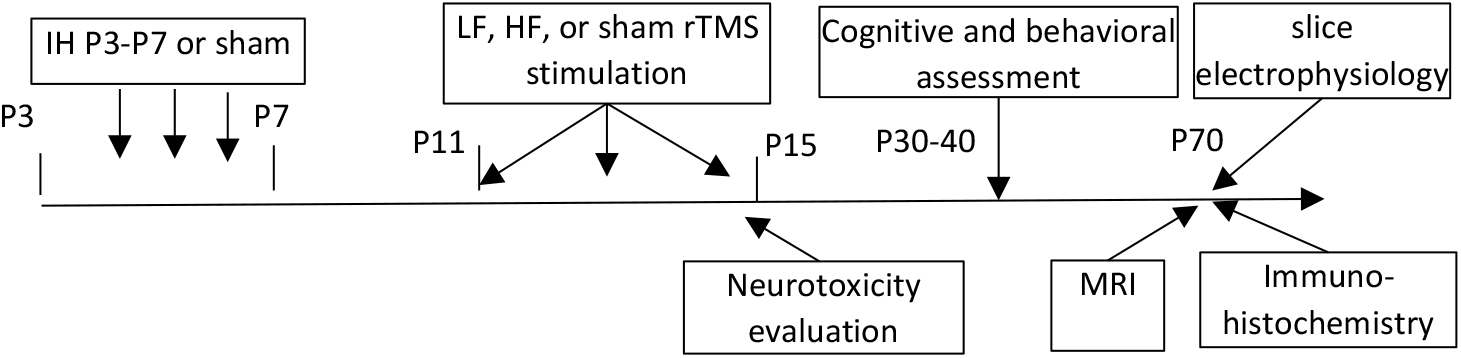
Timeline of the experiments.

### Neonatal intermittent hypoxia

C57BL/6J mice were obtained from the Jackson Laboratory (Bar Harbor, Maine) and bred at the Endeavor Health animal facility. The protocol to induce IH brain injury in a neonatal mouse model, mimicking apnea prematurity in human infants, has been described earlier ^18^. Neonatal mice of both sexes were randomly assigned to the IH or control groups. Animals underwent an intermittent hypoxia protocol involving cycles of hypoxia with a 5% O_2_ / 95% N_2_ gas mixture for 2.5 minutes, followed by 5 minutes of normoxia with room air. During hypoxic exposure, pups were separated from their dams and placed into a 250 mL airtight chamber inside a temperature-controlled neonatal incubator set at 35 °C. The protocol started on postnatal day 3 (P3) and continued for five consecutive days. Airflow was maintained at 4 L/min and alternated between the hypoxic mixture and room air using solenoid valves controlled by a programmable timer. Oxygen concentration and air temperature within the chamber were continuously monitored with calibrated sensors, reaching a nadir of 5% O_2_ within 40 seconds during hypoxic periods. The total number of daily hypoxic episodes gradually decreased over the 5 days (20, 18, 11, 9, and 8) across morning and afternoon sessions. Between sessions, pups were returned to their dams for 4 hours to recover and nurse. Control mice experienced similar handling procedures but remained in room air inside the incubator throughout the experiment. Mortality after neonatal IH was 13.5%. The number of animals used for each outcome measure is provided below.

### Transcranial magnetic stimulation

At P11, mice were randomly assigned to groups receiving repetitive transcranial magnetic stimulation (rTMS). rTMS was applied to non-sedated mouse pups using the Neurosoft Transcranial Magnetic Stimulator (Soterix Medical Inc., Woodbridge, NJ) and a liquid-cooled 58x35 mm figure-of-eight TMS coil, optimized for unilateral stimulation in rodent experiments ^19^. The coil can deliver 2.6 T at 80% power. Stimulation involved a train of pulses over 10 minutes, delivered twice daily with a 4-hour interval between sessions, from P11 to P15 for 5 consecutive days. The groups included Low frequency (1 Hz pulse train, 600 pulses per session), High frequency (10 Hz for 6 seconds, 54 seconds rest, 600 pulses per session), and Sham stimulation (n=8,7), which used the same setup but without actual stimulation. The mouse’s head was immobilized in a custom-printed helmet (Figure 1), and its body was lightly wrapped in gauze to reduce movement. The TMS coil was positioned 5 mm above the bregma to target the bilateral sensorimotor cortex. Stimulation was set to 70% of the device’s maximum power, about 10% below the motor threshold evoked by a single pulse. Motor responses were observed visually through hind limb and tail twitching (Supplementary video 1). Mice in sham control groups were head-immobilized for the same period but did not receive stimulation. To reduce stress, mouse pups were habituated the day before stimulation by handling and gently restraining them for 5 minutes.

**Figure 1.**
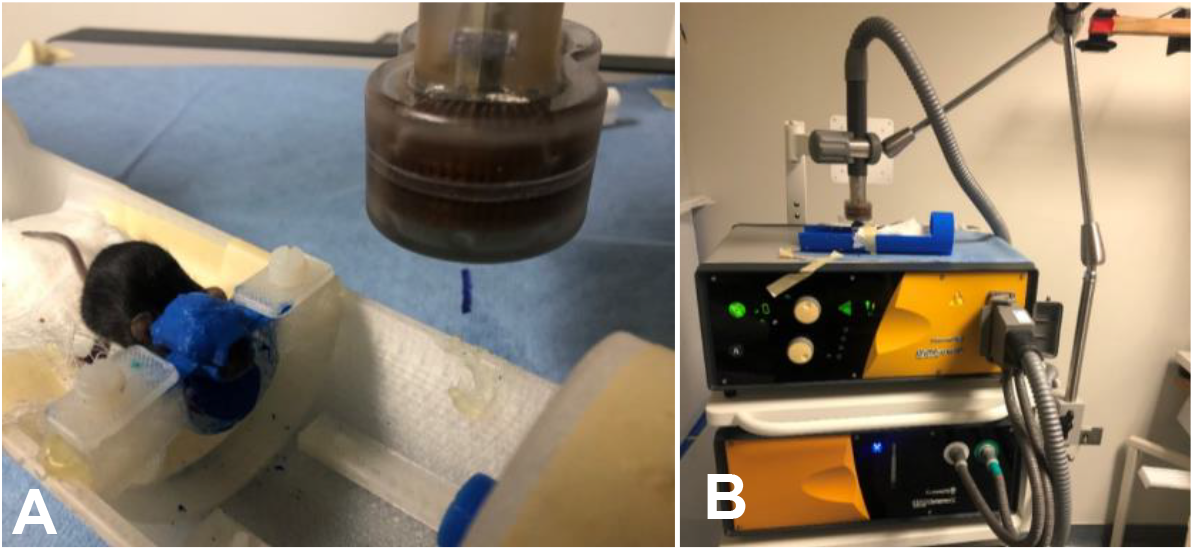
A. Application of transcranial magnetic stimulation on a P12 mouse pup with a dedicated actively cooled rodent 25 mm TMS coil. A custom helmet enables pup head immobilization for rTMS. B. View of the stimulator experimental setup.

### In vivo MRI methods

Ten-week-old mice were sedated with isoflurane (Abbott, IL) inhalation, diluted in air to 5% for induction and 1.5% for maintenance. The animals’ respiratory rate and rectal temperature were monitored with a small-animal physiological monitor (SAII’s Small Animal Instruments, NY, USA). Body temperature was maintained at 35 °C by blowing warm air from a temperature control unit. The animals were placed prone in a cradle and imaged in a 9.4 T Bruker Biospec system (Bruker, Billerica, MA). The receiver coil was a 16 mm diameter, linear surface coil (Doty Scientific, Columbia, SC). The transmitter was a 70 mm quadrature volume coil.

Macromolecular proton fraction (MPF) mapping was performed to assess myelination quantitatively. MPF has been histologically validated as a sensitive myelin marker in the white and gray matter of normal rodent brain ^20^, and experimental models of toxic demyelination ^21^, remyelination ^22^, acute and chronic stroke ^23^, and developmental perturbations ^24^., including the mouse IH model ^18^. 3D MPF maps were generated from three source images (magnetization transfer (MT), proton density, and T1-weighted using a single-point approach with synthetic reference images ^25^. PD- and T1-weighted GRE images were acquired with TR/TE = 16/3 ms and flip angles of 3° and 16°, respectively. MT-weighted images were obtained with TR/TE = 22.5/3 ms and a flip angle of 8°. The off-resonance saturation pulse was applied at an offset frequency of 4.5 kHz with an effective saturation flip angle of 500°. All images were captured in the axial plane with whole-brain coverage and a resolution of 0.15×0.15×0.3 mm^3^. Each image set was acquired with four signal averages. The 3D imaging experiments used a linear phase-encoding order, included 100 dummy scans, employed slab-selective excitation, and collected a fractional (75%) k-space in the slab-selection direction. To correct for field inhomogeneities, 3D B0 and B1 maps were acquired using the dual-TE method (TR/TE1/TE2 = 20/2.9/5.8 ms, α = 8°) and actual flip-angle imaging (TR1/TR2/TE = 13/65/4 ms, α = 60°), respectively ^26^. The total imaging time was 26 minutes. All reconstruction processes were carried out with custom-designed C-language software available at https://www.macromolecularmri.org/.

Magnetization transfer images of mouse brains, obtained as part of an in vivo MPF experiment and showing excellent white/gray matter contrast, were used for automatic structural parcellation using a multi-atlas label fusion method, as described in ^27^. Briefly, individual mouse head images underwent brain extraction, intensity non-uniformity correction, affine registration to atlas images, and label fusion. The publicly available MRMNeAt atlas database was used, which contains 10 individually labeled in vivo mouse brains of the C57BL/6J strain. MPF values in twenty-one white and gray matter structures from the atlas were extracted for each animal.

### Ex vivo field excitatory postsynaptic potential (fEPSP) recordings in motor cortex

Electrophysiological studies were performed in adult mice after behavioral and MRI assessments at P70, with 4 mice per group. Mice were deeply anesthetized using 3% isoflurane and then decapitated. Brains were quickly extracted into ice-cold artificial cerebrospinal fluid (aCSF) composed of (mM): 124 NaCl, 2.5 KCl, 25 NaHCO_3_, 2 CaCl_2_, 2 MgSO_4_, 1.25 NaH_2_PO_4_, 10 HEPES, 10 D-glucose, bubbled with carbogen (95% O2/5% CO2) to maintain a pH of 7.4. All drugs were sourced from Sigma-Aldrich Inc., MO, USA). The osmolarity of this solution was maintained at 300±5 mOsm. Slices containing the motor cortex were cut at 300 µm thickness using a Leica vibrotome (Leica Biosystems). Slices were kept in an incubation chamber filled with room-temperature (21°C) aCSF, bubbled with carbogen, for at least 1 hour before recording. The same aCSF solution was used during recordings and perfused at 3 mL/min at 21°C. After incubation, the slices were transferred to a submerged recording chamber (model RC-24, Warner Instruments, Hamden, CT, USA), where they were perfused at 3 mL/min by gravity. A glass recording electrode filled with aCSF was positioned in the middle of layer six of the M1 cortex. A bipolar stimulation electrode, made of tightly twisted 25 µm diameter platinum/iridium wires, was placed 200-400 µm from the recording site within the molecular layer. The inter-electrode distance was adjusted to prevent pop spikes, which could cause secondary activation via stimulated afferent cells at stimulation intensities ranging from 0 to 700 µA. Input-output curves were created by plotting the amplitude of responses against stimulation intensity. Stimulus pulses of 0.1 ms duration were delivered from the stimulus isolator. Evoking fEPSPs involved applying stimuli once per minute throughout the recordings. Recordings were made using a Multiclamp 700B patch-clamp amplifier (Molecular Devices), with the low-pass filter off to measure the direct current potential relative to the baseline. A potential offset was applied to the pipette before slicing. The amplifier’s high-pass filter was set at 3 kHz to enhance the resolution of the initial slope. Data were sampled at 100 kHz with a Digidata 1442 and analyzed with pClamp 10.1 software (Molecular Devices). Input-output curves were generated by increasing stimulus intensity in 20-μA steps from 0 to 300 μA, while recording fEPSP responses. Paired pulse facilitation (PPF) was tested with two stimuli separated by 70 ms; PPF was expressed as the ratio of the second to the first fEPSP amplitude.

### Behavioral studies

Behavioral testing procedures were conducted in a dedicated quiet room during the daytime when the animals were between 30 and 40 days old. Each apparatus was thoroughly cleaned with 70% ethanol and allowed to air dry for 3 minutes between animals.

Open Field Test: Mice (n = 6 per experimental group per sex) were individually placed in the center of a clear, open-field box (61 x 61 cm) and allowed to explore freely for 20 minutes. Animal movements were tracked and analyzed using ANY-maze software version 7.3 (Stoelting Co., Wood Dale, IL, USA). The software automatically recorded the following parameters of spontaneous locomotor activity: total mobile time, mean speed, total distance traveled, and time spent in the center zone (40 x 40 cm) compared to the peripheral zone. The time spent in the center zone is often used as an indicator of anxiety-like behavior in rodents.

Rotarod Test. Motor coordination and balance were assessed using an accelerating rotarod apparatus (Harvard Apparatus, Holliston, MA, USA) ^28^. Mice (n=5 per sex) were placed on a rotating drum (3 cm diameter) initially set to 4 rpm and accelerated to 40 rpm over 5 minutes. Each animal underwent three trials per session with at least 10 minutes of rest between trials. Latency to fall was recorded, and the best-performing trial was used for analysis. Animals that completed the full duration without falling were assigned the maximum score. All equipment was cleaned between trials to prevent odor cues.

### Cell injury assessment on real-time PCR

A subset of mice (n=4 per group) was euthanized at P15, the final day of the rTMS course. Brains were collected, individually frozen in liquid nitrogen, and stored at -80°C before use. The tissue was homogenized on ice in Qiazol Lysis Reagent (cat. # 79306 Qiagen, Germantown, MD) for total RNA extraction. The NanoDrop (ThermoFisher Scientific, USA) was used to assess RNA quantity and quality. 500 ng of isolated total RNA served as the template for cDNA synthesis using an RT2 First Strand Kit from QIAGEN. cDNA was amplified via PCR with the QuantiTect Sybr Green PCR Kit (Qiagen, Germantown, MD) on an Applied Biosystems QuantStudio 7 Flex real-time PCR instrument. The total reaction volume was 25 μL, containing 1.0 μL of RT product (cDNA). Primers for Caspase-3, IL-1β, IL-6, TNF-α, and GAPDH (Integrated DNA Technologies, Inc., USA) were used at a final concentration of 0.4 μM, with annealing temperatures adjusted accordingly. Gene expression levels were normalized to GAPDH and expressed as fold change using the ΔΔCt method.

### Immunohistochemistry and assessment of OL subpopulations

Mice were euthanized at P70 by transcardial perfusion with saline solution (0.9% NaCl), followed by 4% paraformaldehyde (PFA). Brains were removed and post-fixed in 4% PFA overnight, soaked in 30% sucrose solution for cryoprotection, embedded in OCT, frozen on dry ice, and cryosectioned at 20 μm in the coronal plane.

To identify oligodendrocyte progenitor cells (OPCs) and maturing oligodendrocytes, primary antibodies against NG2 (ab275024, Abcam) and Olig2 (ab9610, Millipore) were used, respectively. Three to five animals per group were analyzed. For each mouse, three coronal sections at the bregma levels of +1 mm, +0.5 mm, and 0 mm were prepared. The sections were hydrated in 0.01 M PBS, blocked in 5% normal goat serum, and incubated with the primary antibodies at 4 °C overnight. Sections were incubated with the appropriate fluorescent secondary antibodies—Alexa Fluor 488 anti-rabbit (ab150077, Abcam) or Alexa Fluor 488 anti-chicken (ab1510169, Abcam)—for 1 hour at room temperature. Subsequently, sections were mounted using DAPI-containing mounting medium (Vectashield, H-1200-10). Negative controls were prepared by omitting the primary antibodies and incubating sections only with the secondary antibodies.

For each animal, three sections were analyzed to estimate the numbers of microglia, astrocytes, oligodendrocytes, and OPCs, using a Nikon E600 microscope and the Cell Counter plugin in ImageJ (https://imagej.net/Downloads). Three counting frames of the corpus callosum and three of the internal capsule were placed on each section at 20× magnification. Cells positive for IBA-1, GFAP, NG2, and Olig2 were counted within a frame measuring 100 × 400 pixels (62 × 251 μm). Cell counts were normalized to the total area of the counted frames. All image acquisition and quantification procedures were performed by an investigator blinded to the experimental groups to ensure unbiased analysis.

### Statistical analysis

A two-way ANOVA was used to test the main effects of rTMS and IH factors on each experimental measure, followed by Bonferroni’s *post hoc* multiple comparison tests. Analysis was performed using GraphPad Prism, version 10.5 (GraphPad Software, La Jolla, CA, USA).

## RESULTS

### rTMS does not induce apoptosis and inflammation in neonatal mice

Analysis of gene expression for the apoptotic marker Caspase-3 and pro-inflammatory cytokines TNF-α, IL-1β, and IL-6 did not reveal significant effects for the main factors IH, rTMS stimulation, and their interactions (F_2,22_=0.47, p= 0.69 for Caspase-3, F_2,22_=0.50, p= 0.61 for IL-1β, F_2,22_=2.50, p= 0.10 for IL-6). The results (Fig. 2) confirm previous findings of little cell death in the neonatal chronic IH model 18 and indicate that the performed rTMS regimen has no apoptotic or pro-inflammatory effects on the neonatal mouse brain. There was a decrease of TNF-α after rTMS in the IH group (rTMS main effect F_2,22_=4.2, p= 0.029).

**Figure 2.**
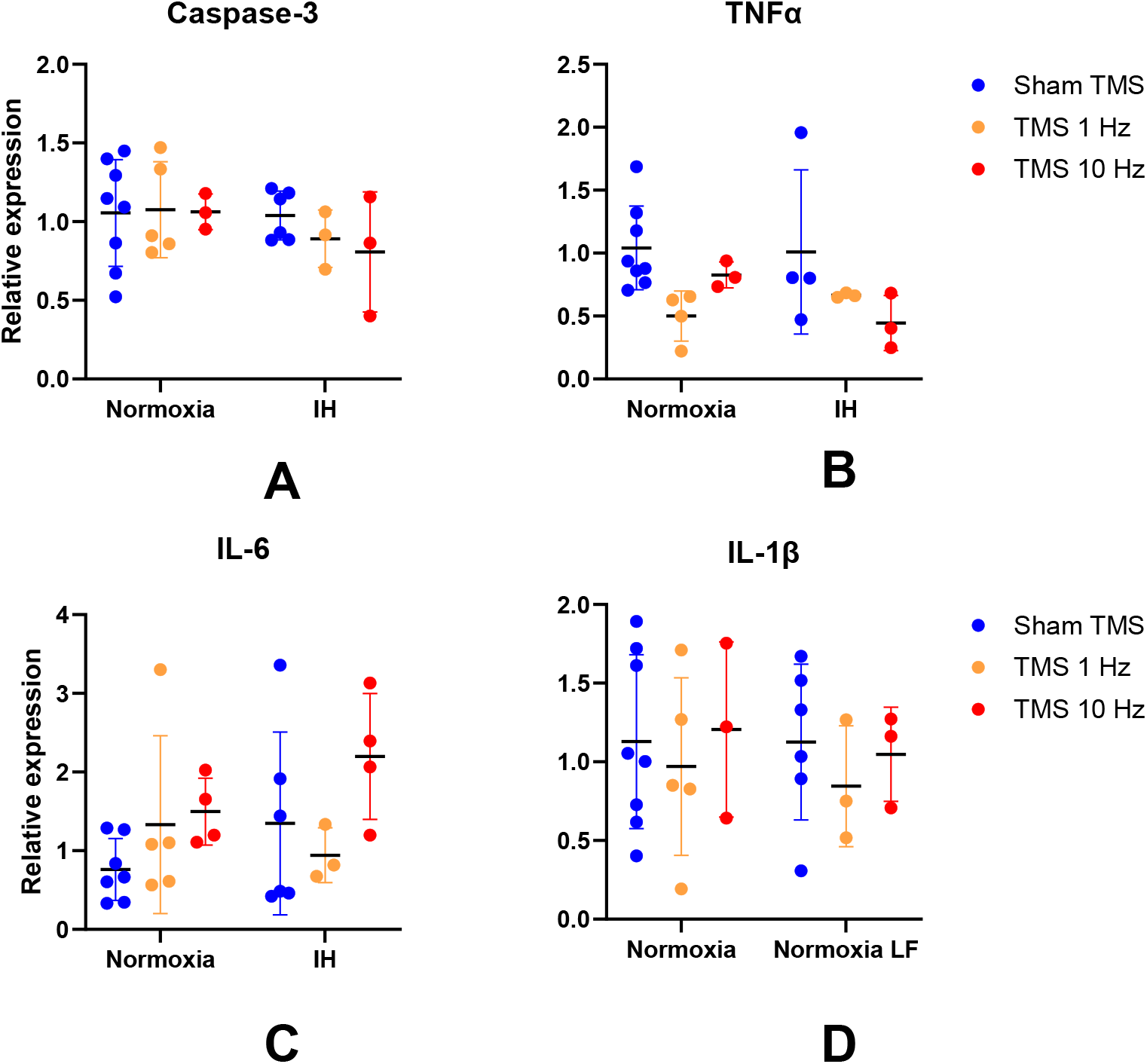
rTMS did not induce apoptosis and inflammation in neonatal Normoxia and IH mice. Relative expression of genes to housekeeping gene GAPDH, determined by RT-PCR, is shown for apoptotic marker caspase-3 (A) and pro-inflammatory markers TNFα (B), IL-1β (C), and IL-6. No significant main effects on IH and rTMS on 2-way ANOVA. Data are shown as mean ± SD.

### HF-rTMS ameliorates hypomyelination after neonatal IH in the corticospinal tract

Representative MPF maps for the sham and HF-rTMS mice are shown in Figure 3. Note a bilateral increase in myelin content in the internal capsule of the stimulated mouse (Figure 3B, arrow).

**Figure 3.**
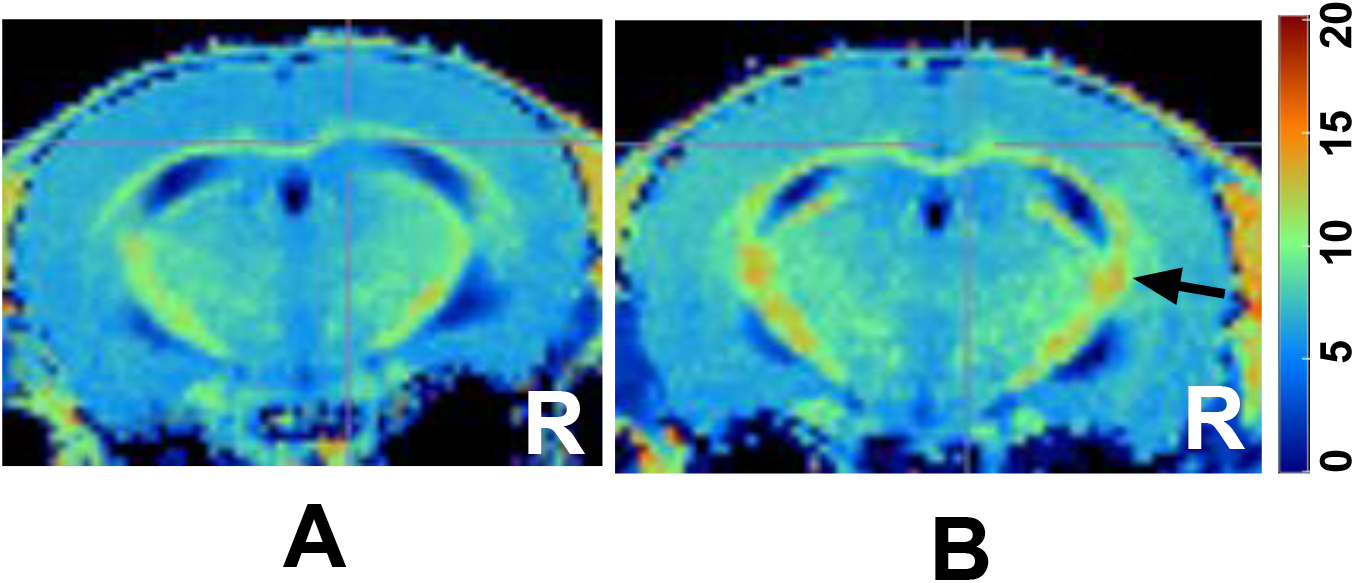
MPF maps in sham (A) and 10 Hz rTMS mouse (B). The arrow points to the internal capsule in the right hemisphere, which was targeted for stimulation. The color bar indicates the macromolecule proton fraction for the pseudo-colored maps in % values.

The main effects of the IH (F (2,38) = 6.42, p=0.0039), rTMS (F (1, 38) = 5.18, p=0.028), and their interaction (F (2,38) = 3.31, p=0.043) were significant in a 2-way ANOVA on MPF in the right internal capsule of the hemisphere targeted by the rTMS stimulation (Figure 4 A). Post-hoc comparisons showed a larger increase in myelination with HF-TMS (p=0.004) and a more minor increase with LF-rTMS (p=0.031) compared to sham stimulation in the IH group. A decrease in myelination after IH and rTMS stimulation (IH (F (2,38) = 3.30, p=0.047) and rTMS (F (1, 38) = 7.10, p=0.012)) was also seen in the internal capsule of the opposite hemisphere (Figure 4 B). Post hoc multiple comparisons revealed a significant difference between the LF-rTMS groups (p=0.046). No significant effects of IH and TMS stimulation were found in the corpus callosum and cerebral cortex (Figure 4C, D).

**Figure 4.**
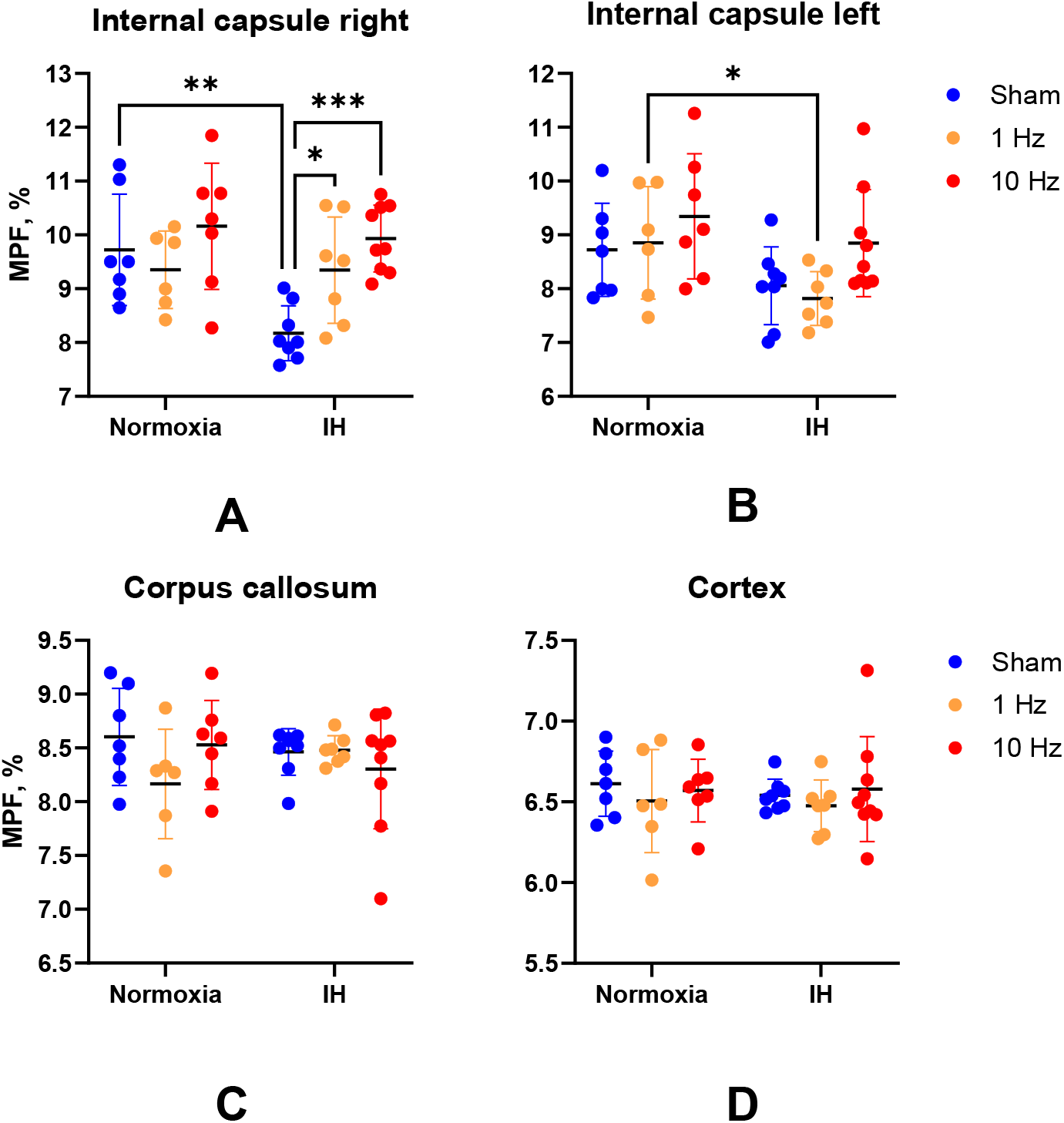
Macromolecule proton fraction group 2-way ANOVA with post-hoc multiple comparisons. A. Internal capsule right (stimulated) and left (B). No significant main effects in IH and rTMS in Corpus callosum (C) and Cortex (D) on 2-way ANOVA. Data are shown as mean ± SD.

### Low-frequency rTMS reduces locomotor hyperactivity after neonatal IH

Female and male mice were analyzed separately in the open field test. After neonatal IH, female mice exhibited hyperactivity, characterized by increased locomotion speed, more time spent mobile, and a greater traveled distance, as shown in Figure 5A (post-hoc comparisons of means p=0.016). However, the overall main effect of IH on traveled distance was not significant in the 2-way ANOVA (F (1, 30) = 1.61, p = 0.21). The main effect of rTMS was significant (F (1, 30) = 4.39, p = 0.044), although the interaction between IH and rTMS was not (F (2,30) = 2.80, p = 0.076). Additionally, there were no significant effects of IH and stimulation on travel distance in male mice.

**Figure 5.**
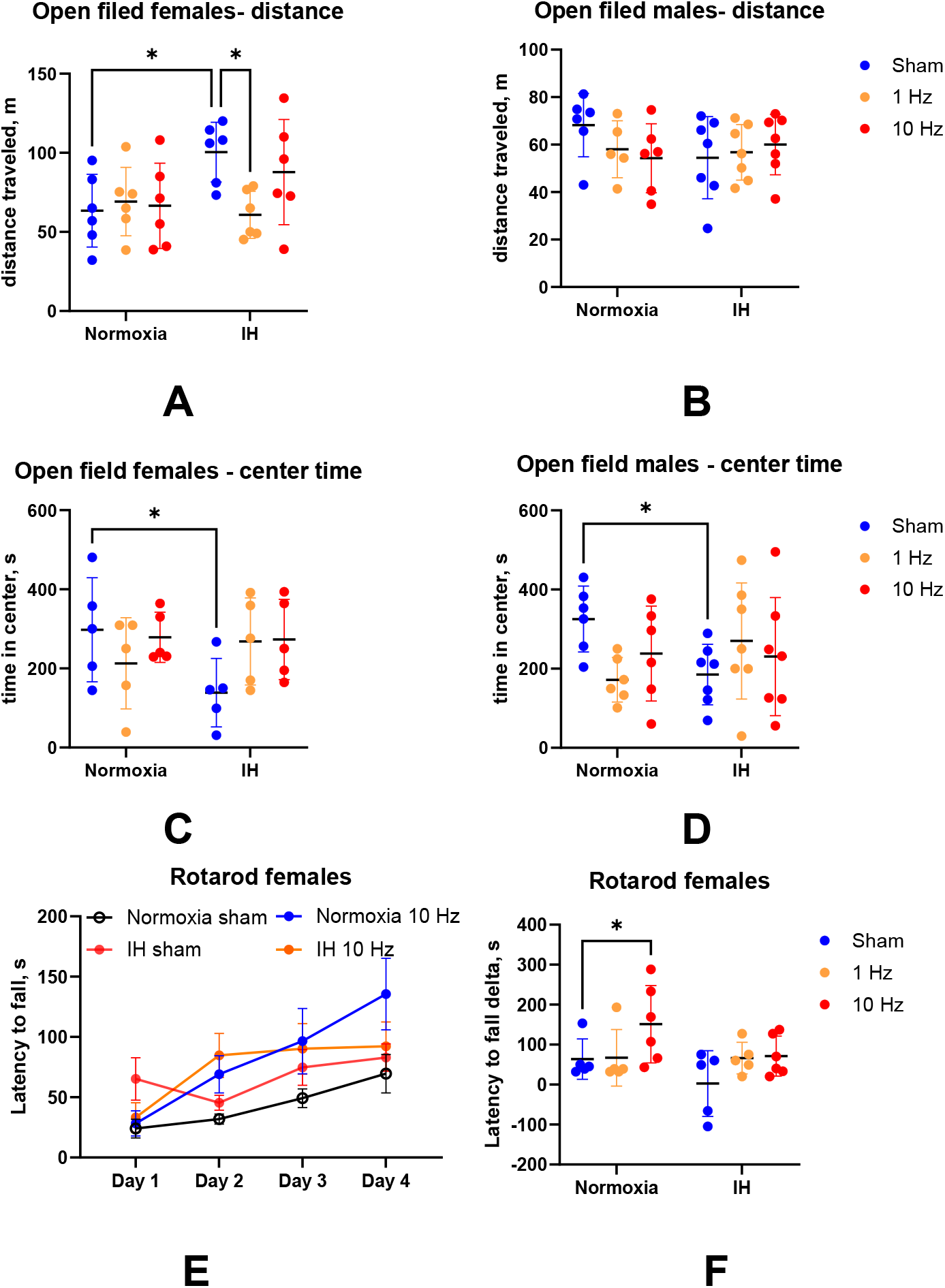
Behavioral tests in mice after neonatal IH and rTMS stimulation on 2-way ANOVA. A, B - Travel distance in open field test for female (A) and male mice(B). C,D – Time spent in the center zone in the open field test for female (C) and male mice (D). Data are shown as mean ± SD. E. Latency to fall on rotarod test from day 1 to day 4 of testing, D. A change of latency to fall from day 1 to day 4. *-p<0.05, post hoc test on 2-way ANOVA

Analysis of time spent in the center area of the open field arena, recorded as an index of anxiety, did not reveal significant main effects for IH or stimulation, but did reveal a significant interaction term in males (F(2,33) = 3.61, p = 0.038). The mean time spent in the center zone was significantly less for the IH sham groups for females. For males (Figure 5C, D, post-hoc p<0.05), but there was no difference with the sham groups after rTMS, indicating anxiolytic effect of rTMS after neonatal IH.

### High-frequency rTMS enhances motor learning in normoxic mice but not after neonatal IH

Latency to fall on the rotarod test increased from day 1 to day 4 in all experimental groups, indicating motor learning in coordination and balance (Figure 5 E, show for females), with significant factors days of test (F(3,99) = 13.413.61, p<0.001) and treatment-by-day of test interaction (F (9.,99) = 2.88, p=0.048) on repeated measures ANOVA. There was no difference in latency to fall on day 1; however, the increase in latency to fall differed by treatment group (Hypoxia factor: F(2,26) = 3.57, p = 0.042 in a 2-way ANOVA). The motor improvement was the most prominent in the Normoxia HF-rTMS group in females relative to the Normoxia sham (post-hoc p=0.045, Figure 5F). No significant difference in the motor learning index was observed between the experimental groups in males on the rotarod test.

### rTMS reduces increased glutamatergic transmission after neonatal IH

The sequence of field responses on stimulation in layer 6 of motor cortex perfused slice preparation, consisting of the short latency volley burst (VB) followed by the first short latency fEPSP and often by the longer latency second fEPSP, was reproducible in all subgroups and observed in both control aCSF and after PTX application (Fig.6A). The VB was insensitive to neurotransmitter receptors blockade and presumed to be presynaptic. The later components, first and second field potentials, were completely blocked by 50 µM CNQX and reflected monosynaptic (latency 4-6 ms) and polysynaptic (latency 6-10 ms) activity, both mediated by glutamate via AMPA receptors (Wallace, Jackson et al. 2014). There were no other components in the field potentials, such as NMDA, kainate, or GABAergic transmission components, after CNQX and PTX block. VBs were removed entirely by blocking axonal transmission with one µM of Tetrodotoxin (TTX). Input-output curves are shown in Figure 6B as mean values of the first field potential amplitude. Pair-pulse facilitation did not differ between the experimental groups (Figure 6B; one-way ANOVA, F(5,18)=0.75, p=0.57), indicating that glutamate release probability was not altered.

**Figure 6.**
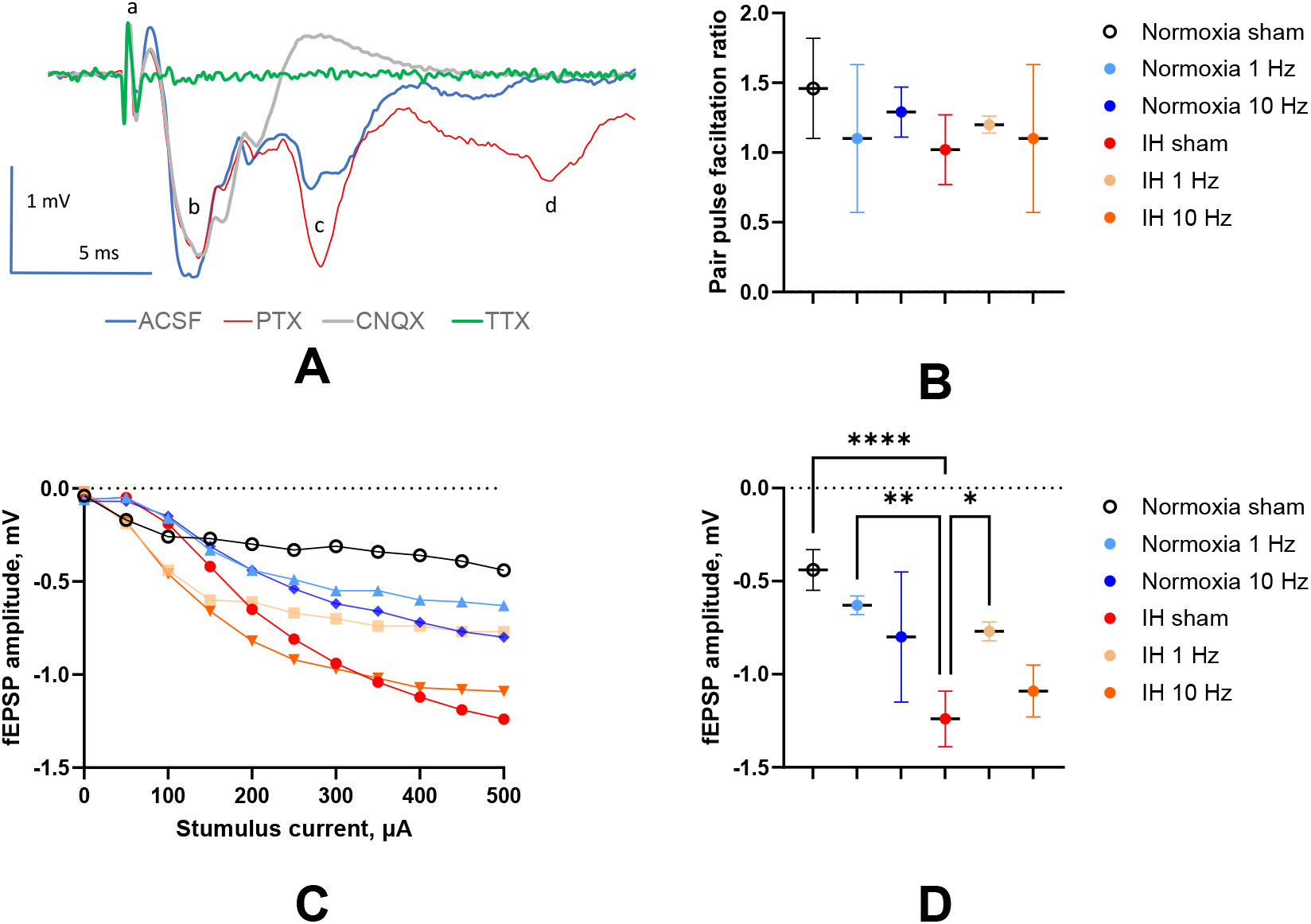
Effect of IH and TMS on post-synaptic field potential in mouse motor cortex slices. A. Examples of fEPSP and waveforms originating at P70. a - truncated stimulation artefact, b – volley burst, c – first monosynaptic fEPSP, d-second polysynaptic fEPSP. The legend below shows inhibitors used to identify the origin of components. B. Pair pulse facilitation ratio was not different between the experimental groups. C. Input-out curves were quantified at stimulus intensity 500 µA with one-way ANOVA (D). Significance of post-hoc comparison is indicated as *-p<0.05, **-p<0.01, * p<0.001

Synaptic excitability varied among the groups and was assessed by measuring the amplitude of the fEPSP at a stimulation intensity of 300 µA on input-output curves (Figure 6D; one-way ANOVA, F(5,18)=11.42, p<0.0001). Post-hoc multiple comparisons showed that synaptic excitability was higher in sham IH animals (Fig. 6D) than in Sham Normoxia (p<0.0001). This increase in excitability after IH was normalized after 1 Hz rTMS treatment (p=0.013) to a level not significantly different from Sham Normoxia.

### rTMS does not affect oligodendrocyte progenitors and total number in normoxic controls and after IH

To investigate whether rTMS treatment could promote the proliferation of immature oligodendrocytes (OPCs), NG2-positive cell counts were performed in the CC and IC in the stimulated (right) and non-stimulated hemispheres. In the CC, significant effects were found for the IH factor (F(1,12) = 8.657, p = 0.0123) and the interaction term (F(2,12) = 4.496, p = 0.0349), whereas the rTMS factor was not significant (F(2,12) = 0.9224, p = 0.4240). In the right internal capsule (IC), no significant effects were observed for IH (F_1,12_= 3.868, p = 0.0728), rTMS (F(2,12) = 3.512, p = 0.0630), or their interaction (F(2,12) = 0.9584, p = 0.4110). Similarly, in the left IC, no significant effects were detected for IH (F(1,12) = 0.1825, p = 0.6760), rTMS (F(2,12) = 0.8185, p = 0.4643), or the interaction term (F(2,12) = 0.0133, p = 0.9868). There was no significant effect of IH and rTMS in the total numbers of oligodendrocytes in the studied white matter tracts.

**Figure 7.**
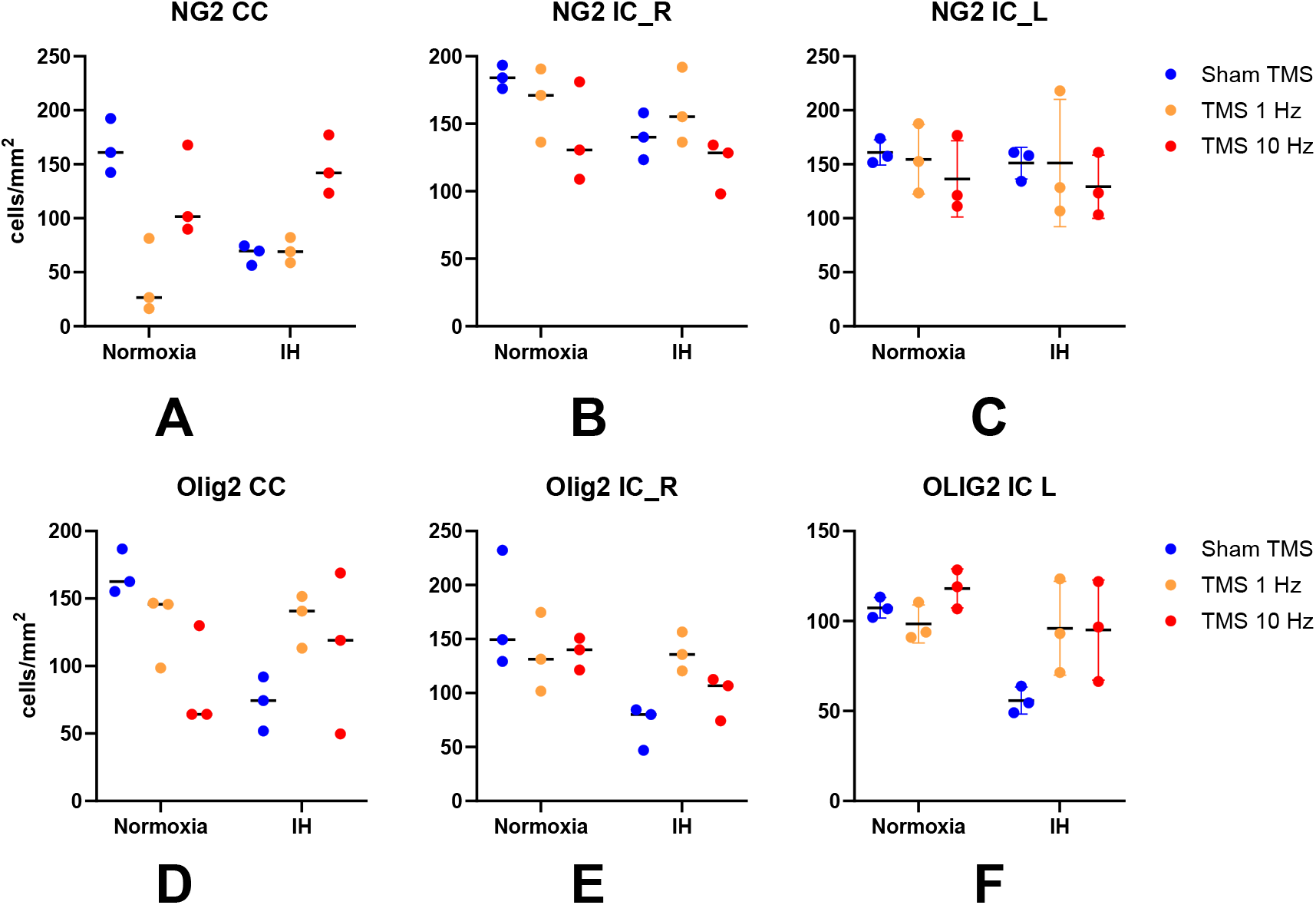
Histological evaluation of proliferation and differentiation in oligodendrocyte lineage. Quantification of NG2-positive cells was performed in the CC (D), right IC (E), and left IC (F). Olig2-positive cell counts were evaluated in the CC (G), right IC (H), and left IC (I). Data are presented as mean ± SEM and were analyzed using two-way ANOVA followed by Bonferroni’s *post hoc* test.

## DISCUSSION

This study utilized a comprehensive approach, combining in vivo MRI, ex vivo electrophysiology, histology, molecular biology, and behavioral assessments in a mouse model of neonatal IH to examine the different effects of LF- and HF-rTMS on cortical excitability and white matter myelination. It found that HF-rTMS effectively improved hypomyelination in the corticospinal tract. Conversely, LF-rTMS decreased locomotor hyperactivity and normalized elevated glutamatergic synaptic excitability following neonatal intermittent hypoxia. These different effects highlight the potential of rTMS as a safe and targeted early intervention to address various neurological outcomes of perinatal hypoxic brain injury.

### Safety and Anti-Inflammatory Aspects of rTMS

A critical prerequisite for any intervention in the vulnerable neonatal population is safety. Our results demonstrate that the applied rTMS regimens, both low- and high-frequency, did not induce apoptosis or an increase in inflammatory cytokines in the neonatal mouse brain. This aligns with previous literature on the high safety margin of NIBS in pediatric settings ^10, 11^. The observed decrease in TNF-α expression in the IH group after rTMS suggests a potential anti-inflammatory effect, which could contribute to its therapeutic effects. Pro-inflammatory signals are known to influence oligodendrocyte lineage development negatively, potentially diverting them towards astrogenesis.

### High-Frequency rTMS Ameliorates Hypomyelination

A key finding of this study is the significant improvement in hypomyelination of the corticospinal tract following HF-rTMS after neonatal IH. As quantified by MPF maps, we observed a notable increase in myelin content in the internal capsule, both in the stimulated hemisphere and, notably, in the contralateral hemisphere. The profound and accelerated rise in MPF with HF-rTMS relative to sham in the IH group for the stimulated hemisphere highlights the potential of this stimulation paradigm to promote myelination, a critical process for efficient axonal conduction and functional motor control ^29^. The bilateral effect, despite unilateral targeting, suggests a broader neuromodulation influence of rTMS in the developing brain, potentially via interhemispheric connections or systemic effects. This finding addresses a critical gap identified in the introduction regarding the unexplored potential of non-invasive brain stimulation to promote myelination early after perinatal brain injury. While MPF changes were significant in the CST, no such effects were observed in the corpus callosum or cerebral cortex, suggesting regional specificity or perhaps a need for different stimulation parameters to affect these areas. This regional specificity could be attributed to the nature of the perinatal injury model (primarily affecting CST) and the targeted stimulation, or it could suggest that myelination dynamics differ across brain regions and are differentially responsive to rTMS.

### Low-Frequency rTMS Modulates Cortical Excitability and Behavior

In contrast to the primary impact of HF-rTMS on myelination, low-frequency rTMS (LF-rTMS) demonstrated distinct therapeutic effects, specifically reducing locomotor hyperactivity and normalizing elevated glutamatergic synaptic excitability. Female IH mice exhibited significant hyperactivity in the open-field test, which LF-rTMS effectively reduced. This behavioral improvement was accompanied by a reduction in abnormally high glutamatergic synaptic excitability in motor cortex slices from IH animals treated with LF-rTMS. This normalization of synaptic function supports our hypothesis that LF-rTMS can counteract the excessive glutamatergic transmission and subsequent occlusion of synaptic plasticity observed after neonatal IH ^18, 30^. The anxiolytic effect of rTMS observed in both female and male mice further underscores its broader neurobehavioral benefits. This highlights LF-rTMS as a promising strategy for restoring the excitatory-inhibitory balance, a crucial aspect of normal brain development and function.

### Differential Effects and Therapeutic Implications

The most striking implication of our findings is the apparent differential effect of low- and high-frequency rTMS. HF-rTMS primarily enhanced myelination in the CST, crucial for motor control, while LF-rTMS targeted glutamatergic excitotoxicity and behavioral hyperactivity. This suggests that specific rTMS parameters may be tailored to address distinct pathological aspects of perinatal brain injury. Such precision could overcome limitations of pharmacological approaches, which often entail broad, long-term global inhibition with potentially disruptive consequences on the developing brain. The non-invasive nature and demonstrated safety profile of rTMS make it an attractive alternative for early intervention in a vulnerable population where timing and targeted therapies are critical for preventing the formation of pathological circuits and improving long-term outcomes.

### Limitations and Future Directions

Despite these promising results, this study has several limitations. First, it was conducted in a mouse model, so caution is necessary when applying findings directly to human infants. While mouse models offer valuable insights into underlying mechanisms, there are species-specific differences in brain development and responses to rTMS. Second, our exploration of the molecular mechanisms behind the observed effects is preliminary. Future research should explore in greater detail the cellular and molecular pathways activated by different rTMS frequencies, particularly how they affect oligodendrocyte progenitor cell differentiation, maturation, and myelination, as well as the specific mechanisms underlying synaptic plasticity modulation. Studying inflammatory markers, such as specific pro- and anti-inflammatory cytokines, could also help clarify their role in rTMS-mediated improvement of hypomyelination, as hypothesized.

Furthermore, although we observed positive effects on myelination and synaptic excitability, long-term follow-up studies beyond P70 are needed to determine if these benefits last and how they impact more complex cognitive and motor functions. Future research should also investigate the best timing, duration, and intensity of rTMS treatments, as well as the potential advantages of combining low- and high-frequency protocols to improve therapeutic results. Ultimately, these preclinical findings offer an important foundation for future translational research and clinical trials exploring rTMS as an early, targeted intervention to improve neurodevelopmental outcomes in infants with perinatal hypoxic brain injury.

### Conclusion

Perinatal hypoxic brain injury presents a major clinical challenge, often leading to complex neurodevelopmental disorders such as cerebral palsy and cognitive impairments. Current treatments are mostly palliative, highlighting the urgent need for early interventions that target the underlying pathophysiology. Our findings offer new insights into the therapeutic potential of rTMS as a safe and effective neuromodulation approach for reducing key pathological features of perinatal hypoxic brain injury, including cortical hyperexcitability and white matter myelination issues.

## Acknowledgments

This study was funded by NIH grants 1R01NS119251-01A1

## Notes

Conflict of interest statement: “The authors declare no competing financial interests.”

### Competing Interest Statement

The authors have declared no competing interest.

### Summary of Updates

Added histological analysis Figure 7; author affiliation is updated

